# Integrated optical device for Structured Illumination Microscopy

**DOI:** 10.1101/2022.04.22.489094

**Authors:** Matteo Calvarese, Petra Paiè, Alessia Candeo, Gianmaria Calisesi, Francesco Ceccarelli, Gianluca Valentini, Roberto Osellame, Hai Gong, Mark A. Neil, Francesca Bragheri, Andrea Bassi

## Abstract

Structured Illumination Microscopy (SIM) is a key technology for high resolution and super-resolution imaging of biological cells and molecules. The spread of portable and easy-to-align SIM systems requires the development of novel methods to generate a light pattern and to shift it across the field of view of the microscope. Here we show a miniaturized chip that incorporates optical waveguides, splitters, and phase shifters, to generate a 2D structured illumination pattern suitable for SIM microscopy. The chip creates three point-sources, coherent and controlled in phase, without the need for further alignment. Placed in the pupil of a microscope’s objective, the three sources generate a hexagonal illumination pattern on the sample, which is spatially translated thanks to thermal phase shifters. We validate and use the chip, upgrading a commercial inverted fluorescence microscope to a SIM setup and we image biological sample slides, extending the resolution of the microscope.

## 1. Introduction

Structured Illumination microscopy is an optical technique that is becoming of widespread use for imaging at high resolution and beyond the classical diffraction limit [1]. Initially used for obtaining optical sectioning [2], SIM is now commonly and commercially used for super-resolution microscopy. Structured illumination microscopy can double the resolution of a diffraction limited microscope [3], if it is implemented with linear excitation, where the fluorescence emission is proportional to the excitation intensity. It can further increase the resolution down to the tens of nanometer scale [4], if operated in a nonlinear regime. Recently, structured illumination has opened new possibilities in three-dimensional localization microscopy [5], light sheet microscopy [6] and far field optical nanoscopy [7], among others.

The generation of a coherent SIM patterns is commonly achieved by interference of two or more laser beams on the sample. In an epifluorescence microscope, a sinusoidal pattern can be generated by focusing two coherent laser beams into the pupil of the microscope. By changing the relative phase of beams, the pattern moves (spatially shifts) through the field of view. In its original implementation [3], a diffraction grating created the two coherent beams, and a mechanical translator (and rotator) was used to change the pattern spatial phase and angle. In recent SIM applications the beam is shaped with a spatial light modulator [8,9]. This approach has the advantage to allow the creation of complex modulation patterns, with high phase precision, moreover the spatial light modulator can be further used to measure the microscope aberrations [10]. However, even in modular implementation [11] a bulk optical setup for SIM pattern generation has the disadvantage to be cumbersome and to require day-by-day adjustments. Recently, a few methods to generate a SIM pattern, based on optical fibres and (temporal) phase modulation have been proposed [12, 13], but a fully integrated optical system that does not require free space elements is still lacking.

Here we present an integrated device for patterned illumination, that can be used as a light source for SIM microscopy. The device consists of a miniaturized glass chip that incorporates optical waveguides, beam splitters and thermal phase shifters. The light coupled into the input waveguide of the chip, is split into three waveguides at its output. The relative phases of the three waveguides are controlled by a voltage applied to the thermal shifters. Three optical fibres are connected to the waveguides at the output of the chip and their other ends are placed on a glass ferrule, at the vertices of an equilateral triangle. These three-point sources, azimuthally polarized, interfere at the sample, producing a hexagonal intensity pattern, suitable for SIM microscopy [14, 15], specifically in its hexSIM implementation [9]. By controlling the phase of the three waveguides, we shift the patterns laterally on the sample, acquiring the images required for SIM illumination, demodulation and super-resolved reconstruction. The glass chip, fabricated by femtosecond laser micromachining, integrates the components required for shaping the three sources, maintaining the desired polarization while being able to precisely control the phase.

## 2. Results and discussion

### 2.1 Chip layout

To guarantee a rapid phase shift dynamics the splitting is obtained with a cascade of integrated couplers. The optical circuit (Fig. 1 a,b) is composed of two integrated couplers, which act as beam splitters and distribute laser light among the three output waveguides, according to their coupling ratio. Laser light is coupled into one waveguide through a polarization maintaining fiber (PM). The first splitter is designed with a coupling ratio of 33%, one third of the initial power continues travelling in the first arm, while the remaining is guided to the second splitter. The latter is designed as a 3dB coupler, with a coupling ratio of 50%. With such power splitting, the initial power is ideally equally distributed among the three output ports. In our device we found that the power ratio at the three outputs was 33%, 36% and 30% (Fig. 1c).

**Fig. 1.**
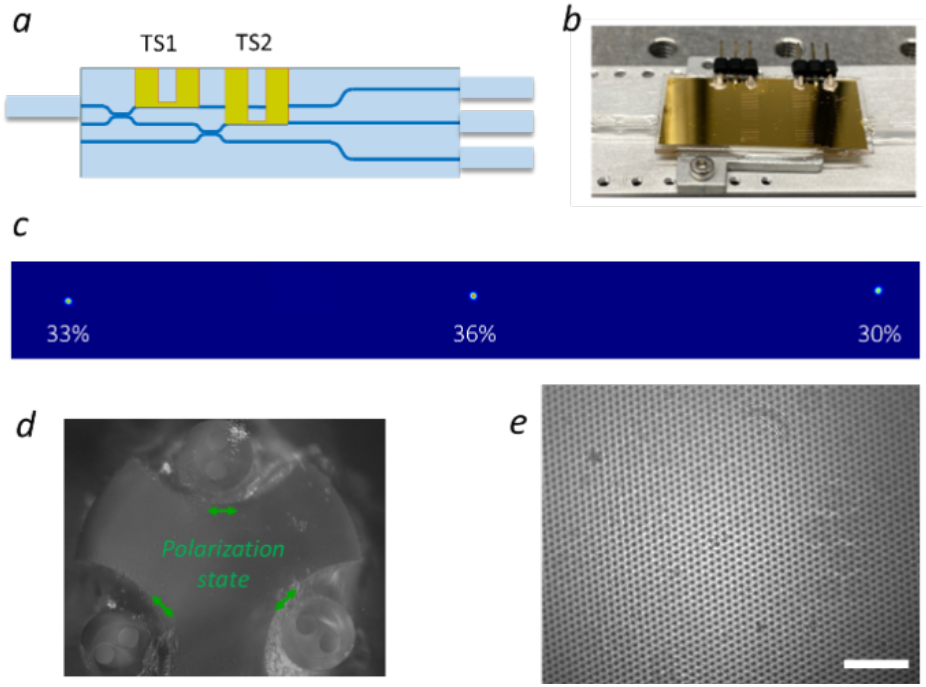
Chip and ferrule layout. (a) Schematics of the integrated optical circuit with two metal thermal phase shifters TS1 and TS2. (b) Image of the real device with pins for electrical connections. (c) The three chip output ports imaged on a camera showing the distribution of power. (d) Image of the actual ferrule, the two small circles in the fibre facet indicates one of the two perpendicular axis of the PM fibres that maintain the polarization. In green the expected polarization state when the input polarization is coupled perpendicular to the circles. (e) Hexagonal pattern formed by focusing the three point sources from the ferrule plane to a camera, using a 50 mm lens. Scale bar is 0.5 mm

The optical circuit is integrated with two thermal shifters (TS1 and TS2 in Fig. 1a) to control the relative phase of the guided beams [16]. The shifters create a phase difference *φ*_*ij*_ between each pair of guided beams *i,j* and they are constituted of a thin metal resistor which is placed on top of one arm of each beam splitter. An applied voltage across them, causes a power dissipation into the chip substrate, and a consequent refractive index difference between the beam paths by thermo-optic effect. This gives a phase shift between two adjacent beams. Using two thermal phase shifters is enough to control the relative phases of the three beams.

The three outputs of the chip are coupled and glued to a PM fibre array, which ends into a specifically designed ferrule. The three fibres tips are arranged in a triangular configuration and their optical axis is aligned before gluing, so to place them in azimuthal polarization configuration (Fig. 1d).

The fabrication of both the active photonic chip and ferrule is based on femtosecond laser micromachining (see Materials and Methods).

### 2.2 Structured illumination microscope

The three-points at the ferrule plane act as the source for the pattern. Thanks to the Fourier transforming properties of lenses, imaging the three coherent points in the back focal plane of a lens, produces a hexagonal intensity pattern in its front focal plane [14]. We initially characterized the pattern placing the ferrule in front of a plano-convex 50 mm lens. This configuration creates an interference pattern which can be collected by a camera placed in the focal plane of the lens (Fig. 1e). As expected, the pattern shows hexagonal symmetry with clear zeros.

We then aimed to create an image of the three point-sources in the microscope’s pupil, so to generate the hexagonal pattern in the object plane. In a commercial inverted microscope this is possible by adding a lens in front of the ferrule, that in combination with the tube lens (TL) of the microscope forms a magnified image of the point sources. We selected this lens to be a 5X microscope objective (Mitutoyo, 0.14 NA). To have more flexibility on the alignment of the illumination and detection path, we added a telescope at the output port of the microscope, composed of two 150 mm lenses (f1 and f2 in Fig. 2).

**Fig. 2.**
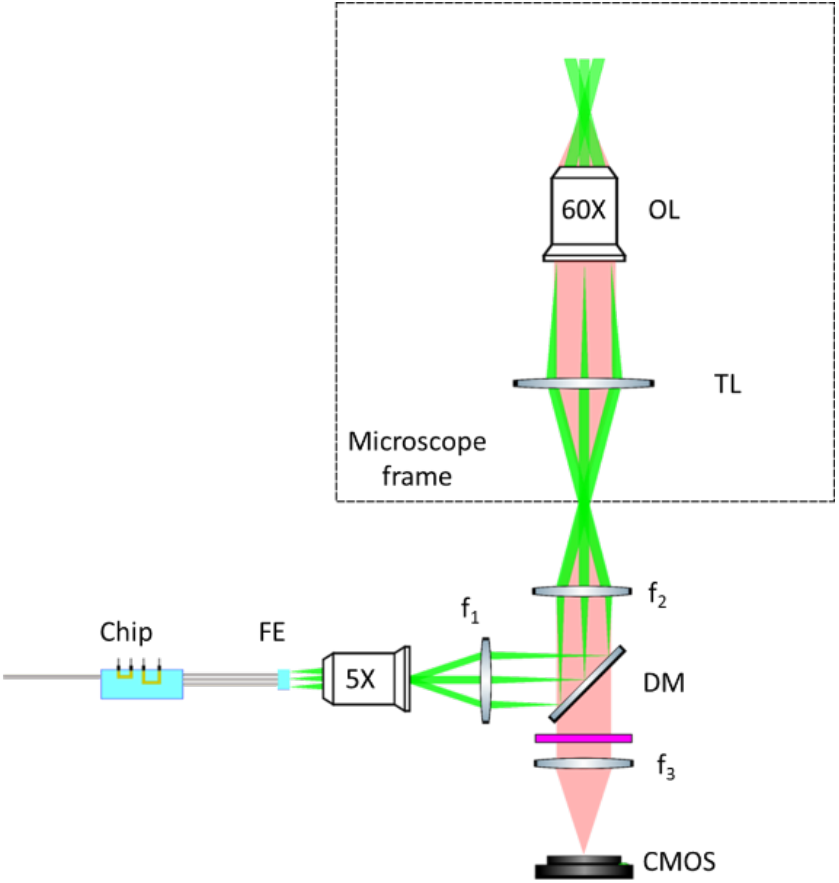
Structured illumination microscope. Green beams represent the excitation while pale red the emitted fluorescence. The dashed line delineates the inverted microscope frame. The chip is connected with polarization maintaining fibers to the ferrule (FE). A low NA objective (5X), together with the first lens (f_1_=150 mm) image the three fibre facets at the ferrule plane onto the dichroic mirror (DM). The second lens (f_2_=150 mm) with the tube lens (TL) of the inverted microscope bring this image to the back focal plane of the microscope objective. The microscope objective (60X) creates the pattern on the sample plane of the microscope. The emitted light is collected by the objective and imaged at the output port of microscope where a relay system formed by lens 2 and lens 3 (f_3_ = 150 mm), creates a secondary image at the CMOS camera. A bandpass filter (BF) eliminates residual excitation light from the detection path.

The three point-sources defined a triangle inscribed in a circle with radius 0.25 mm at the ferrule plane and the circle was magnified to a 1.25 mm radius at the back focal plane of the objective. The pattern period at the sample plane depends on the illumination objective used to form the pattern, its value is affected both by the numerical aperture and the physical size of the pupil. We can in fact define a filling factor FF, as the ratio between the radius of the circle containing the three points in the back focal plane and the pupil radius. The higher this value is, the closer the pattern period gets to the theoretical maximum resolution of the objective.

We used an Olympus 60X oil-immersion objective (1.4 NA). In this configuration, we have obtained a filling factor of ∼0.5, which corresponds to a pattern period of ∼450nm. The filling factor determines the resolution improvement of the SIM reconstruction, as for hexSIM the resolution can be estimated as 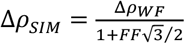 where Δ*ρ*_*WF*_ is the widefield resolution, limited by the microscope objective lens. In the best case the filling factor is 1 and the resolution improvement is 1.87. We tested a filling factor of 0.5, in order to have a clearly visible pattern at the image plane and to demonstrate that the technique is able to generate and move the pattern precisely. Considering that the waist of the beam at the fibres plane is 2.7 μm, at the pupil the waist of each of the sources is 13.5 μm, giving rise to a modulation area with a diameter of ∼ 80 μm (FWHM), at the object.

Figure 3a shows a fluorescent cell (from Invitrogen FluoCells #1 slides labelled with MitoTracker Red), illuminated with the pattern. The hexagonal lattice is superimposed to the fluorescence signal. We proved that a controlled shift of the pattern on the cell is possible, by applying different consecutive voltages to the thermal shifters, and recording the corresponding images. Figures 3b and Fig. 3c show a subsection of the cell, acquired with two different values of phase shifts. In the line plot of Fig. 3d, the pattern translation in the sample plane is visible. This measurement indicates that the chip allows the generation of a SIM pattern on the object plane of the microscope that can be spatially translated in the sample plane by changing the relative phase of the output beams.

**Fig. 3.**
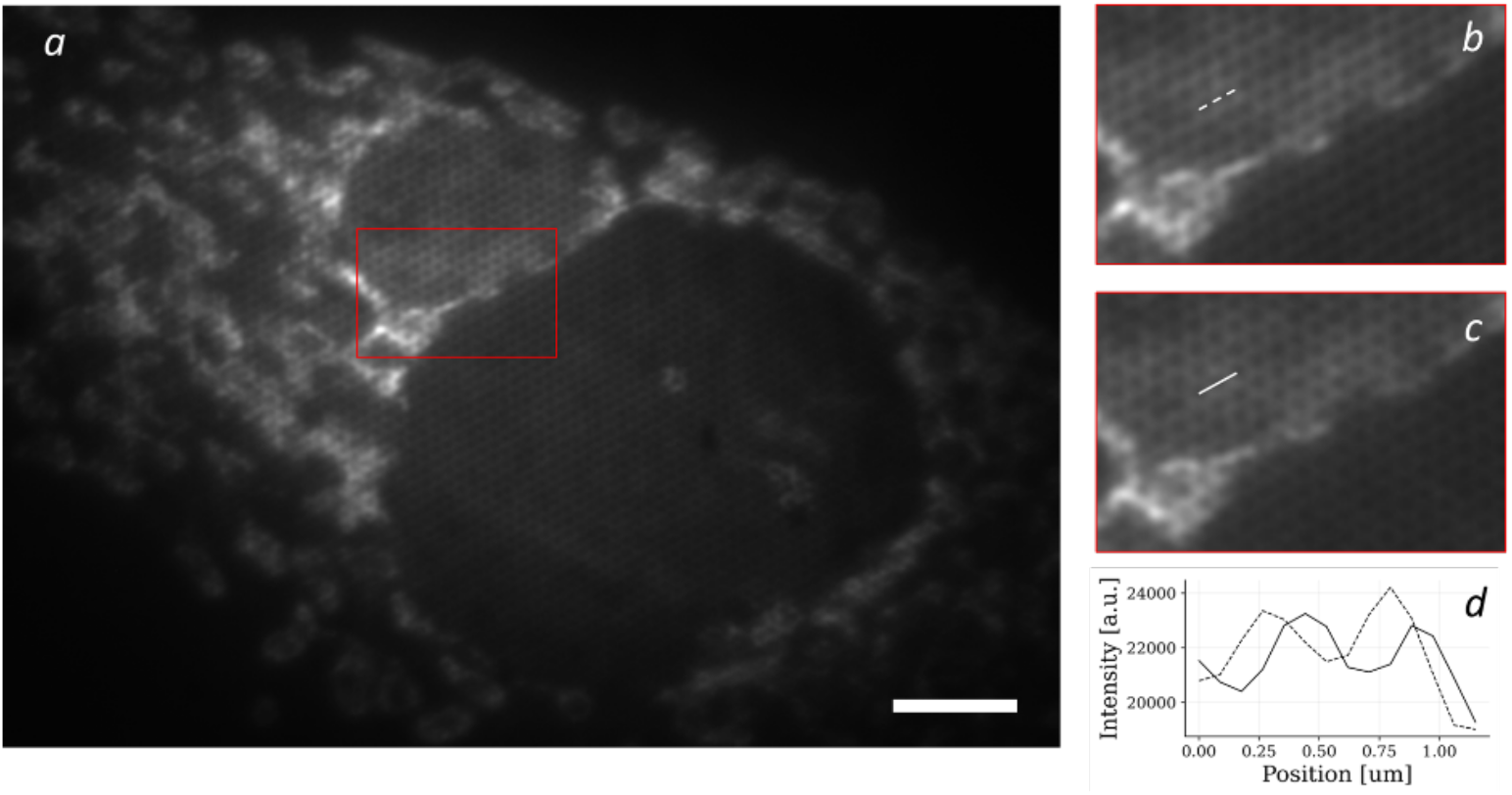
Illumination pattern formation and translation. (a) Illumination pattern formed on a fluorescent cell with a 60X/1.4NA oil immersion Olympus objective. The pattern period is around 450 nm. Scale bar is 5 μm. (b) and (c) zoomed area from panel a acquired with different voltages applied to the thermal shifters. (d) Line plot from the white lines of panel b (dotted) and c (solid). The line plot shows the effective translation of the illumination pattern.

### 2.3 Phase shift and calibration

The hexagonal pattern with active phase control allows one to perform demodulation and to reconstruct super-resolved images acquiring N=7 images with a well precise combination of relative phases between the three laser beams [9]. One advantage of hexSIM consists in having a number of images for reconstruction smaller than the number of images required for standard SIM (i.e. 9 acquisitions corresponding to 3 spatial phases and 3 angles). To implement the N desired phase shifts (see Materials and Methods), we first performed a calibration of the thermal shifters’ response, to assess the voltage-to-phase shift characteristics and find the correct values of input signals. Voltage calibration was performed in a pre-assembling step of the chip, before the output PM fibres were glued. Light was coupled into the chip input and the far field interference pattern of two beams at a time is collected by a camera, while voltage is scanned in a suitable range (0-9V). From post processing of the acquired images, a characteristics curve is obtained, one for each thermal shifter (Fig. 4a).

**Fig. 4.**
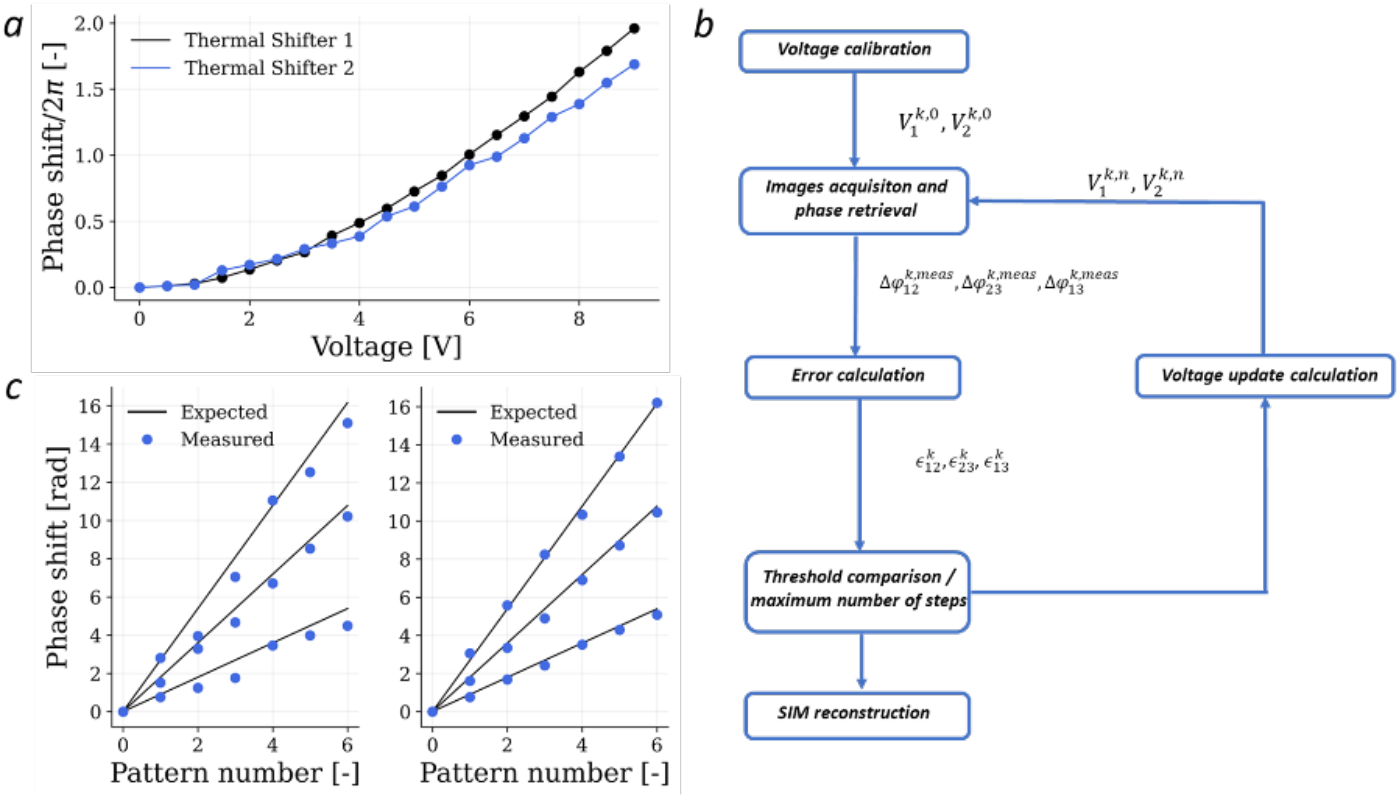
Phase shift calibration. (a) Pre-assembly calibration. The phase shift of two beams is measured while changing the applied voltage to each thermal phase shifter and a voltage-to-phase characteristics is retrieved. (b) Self-calibration block diagram. (c) Comparison between measured (blue dots) and expected (black lines) phase shifts between the three beams before and after 10 cycles of calibration process.

Nevertheless, the precision of this calibration method is not ideal. The reliability of the analogue voltage source, the manual change of the voltage levels and the following gluing process are some of the factors that may affect the final precision of the phase shifting process. Inaccurate phase shifts may result in artefact in the reconstructed image. To circumvent this problem, we implemented a self-calibration method, that exploits an iterative process to calibrate the voltage signals. Initially, images are acquired using the voltage signals found in the standard/analogue calibration, then the phases are calculated from the N acquired images using autocorrelation-based methods (see Material and Methods). The error *ϵ*_*ij*_ between the expected phases and the measured values, for each pattern, is computed as:

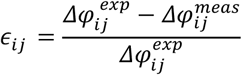

Here the indexes *i* an *j* indicate a generic couple of beams, thus for each pattern we have 3 error values (i.e. *ϵ*_12_, *ϵ*_13_, *ϵ*_23_).

Depending on the value of the errors, a correction to the two input voltages is applied considering the effect of each thermal shifter to the relative phase shift (see Materials and Methods):

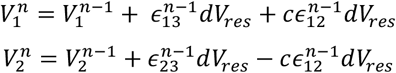

where *n* is the iteration number, *dV*_*res*_ is a voltage step that is set (typically at 0.3V) at the beginning of the calibration process and it indicates the maximum voltage increase for each iteration, *c* is a parameter that takes into account for possible asymmetries in the system. Ideally, only the error on *Δφ*_13_ and *Δφ*_23_ would be enough to set the new voltage values, because *Δφ*_12_ is the difference between them. However, possible nonidealities make it convenient to introduce an additional degree of freedom for the reduction of *ϵ*_12_. The relation between the voltage and the phase is described in the Materials and Methods. A value of c =0.5 was chosen empirically from the resulting phase reconstruction and typically used in our calibration. Its impact on the phases is however limited, so a precise estimate is not necessary.

The process is repeated until the error meets a suitable threshold or a maximum number of iterations M is reached. It is worth noting that, during the process, the region of interest for the autocorrelation analysis is kept the same, so that phase calculation is consistent within consecutive iterations.

Figure 4c shows the comparison between the measured phase shifts for each pattern before and after the calibration process (with M=10). We observe that the phases applied after the calibration are in good agreement with the theoretical ones (Fig 4c, right panel). The average root-mean-square error among the three sets of phases was 0.68 rad (normalised RMS of 16.5%) before optimization and, after 10 cycles of calibration, it was reduced to 0.26 rad (normalised RMS of 5.7%). Once the calibration process is completed, the new voltage signals can be used for the SIM measurement, acquiring N images at the desired phase.

### 2.4 SIM imaging of biological slides

We imaged a cell slide (FluoCells Prepared Slides #1) using a 60x/1.4NA oil immersion objective. The 532 nm laser excites the MitoTracker Red CMXRos that was used to stain the mitochondria of the cells. The pattern created on the samples had a period of 380 nm with a filling factor of 0.57. After the self-calibration process, N=7 images with the correct phases were acquired and the reconstruction process was performed (see Materials and Methods). Fig. 5 compares a widefield image with the reconstruction obtained with SIM. As expected, the reconstructed SIM image shows an improved resolution when imaging the mitochondrial membranes. This improvement is particularly clear by taking a line profile of the filament (Fig. 5c,d). Using decorrelation analysis [17] we estimate a 1.5-fold resolution improvement, in line with the used pupil fill factor. It is worth noting that the reconstruction algorithm reduces the background in the image, significantly improving its contrast, relative to the widefield case. These measurements demonstrate that our device can be used as an add-on system to a standard fluorescence microscope, to implement structured illumination microscopy. We showed the HexSIM configuration, with a pattern formed by a 3-beam interference, but in principle several waveguides could be exploited to be used in N-beams SIM.

**Fig. 5.**
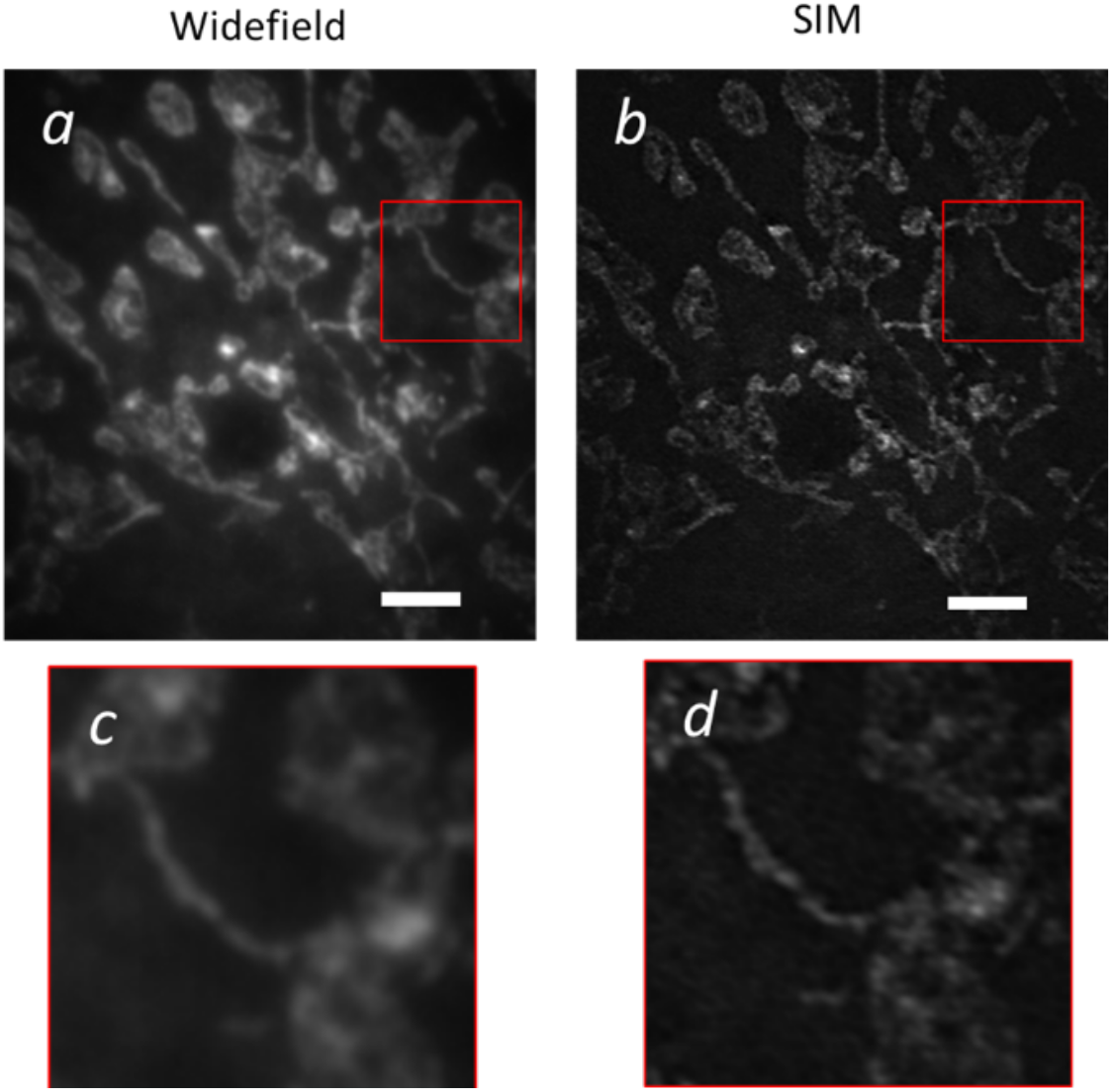
SIM reconstruction. (a) Widefield image of a MitoTracker Red CMXRos stained cell. The mitochondria are visible in the picture. (b) SIM reconstruction of the same region of interest, after the acquisition of 7 phase shifted images and the processing algorithm. The zoom insets show the resolution improvement between widefield (c) and SIM (d). Scale bar is 3μm.

## 3. Materials and methods

### 3.1 Illumination and detection optical system

The system starts with a laser (RGB Photonics, 532 nm, 200 mW) coupled into a polarization maintaining fibre (PM fibre). The fibre is coupled to the chip and fixed at the input port with a UV curing glue (Optical Adhesive 133 by Norland). The output of the waveguides is coupled and fixed to the PM fibre array and the three fibers are then placed on the triangular ferrule. The typical power at the entrance of the chip was 50 mW and the power on each point source was 2.5 mW. The main source of losses are the fibre connections at the input and output facets of the chip, giving an overall transmission efficiency of about 15%.

A 5x microscope objective, together with two 150 mm lenses (Fig. 2) form the hexagonal pattern at the entrance of the microscope port of a Leica DM IRBE inverted microscope. This pattern is brought to the object plane by the tube lens and the microscope objective lens. A dichroic mirror (at 560 nm) is used to reflect the illumination wavelength towards the microscope objective and to transmit the emitted light. Its surface is a conjugate plane with the pupil of the microscope, to facilitate the pattern alignment.

The detection path of the microscope includes a relay system with a shared lens (f_1_) and a second lens in front of the camera (f_3_ = 200 mm). A CMOS camera (FLIR Grosshoper) is used for image acquisition and a bandpass filter (590 ± 15 nm) is used to reduce the residual excitation light.

Phase control is carried out by applying two analogue signals of a National Instruments DAQ board, which provide the input voltages to the electrical pins on the chip. A python code was developed, within the Scope Foundry (http://www.scopefoundry.org/) framework, to control the board, acquire the images and process the data.

### 3.2 Integrated photonic circuit fabrication

The fabrication of the photonic chip consists in different steps. Firstly, the integrated optical circuit was written by FLM in a borosilicate glass substrate (Eagle XG, Corning), using a Yb:KYW cavity-dumped mode-locked laser oscillator, with 300 fs long pulses at 1030 nm, and a repetition rate of 1 MHz. An initial characterization of the waveguide losses allowed us to determine the optimal irradiation parameters. We used a 50x, 0.65 NA as focusing objective, an average laser power of 230 mW, a scan velocity of 40 mm/s and a multiscan approach based on 12 overlapping irradiations. Secondly, a previously optimized thermal annealing process, with a peak temperature of 750 °C [18], followed the irradiation. With this process, we obtained single mode waveguides, with propagation losses of about 0.3 dB/cm and symmetric mode dimensions of about 2.7×2.8 μm^2^ for 532 nm wavelength.

To create a balanced three-beam splitter, we fabricated 3dB directional couplers and we characterized them to find the relationship between coupling ratio and interaction length. The arms distance within the interaction region was fixed to 5 μm. With this characterization, we found the optimal interaction lengths for both the splitters, and we obtained a rather balanced device, with a power ratio at the three outputs of 33%, 36% and 30% (Fig 1c). The interaction length was 0.75 mm for the first splitter and 0.483 mm for the second.

### 3.3 Thermal shifter fabrication and properties

We designed the coupler geometry and the thermal phase shifters based on a previous work [16], where the temporal dynamics, as well as the power consumption of a similar device, were investigated. To have a short temporal response and a low dissipated power, we have fabricated the splitters at a distance of 15 μm from the glass surface and with a distance of 30 μm between the arms. At the output ports, the waveguides are moved apart to 127 μm, to match the spacing of the PM fibres. To create the thermal phase shifters, a metal deposition process was carried out. First, a 2 nm layer of chromium was deposited over the glass substrate to enhance gold-to-glass adhesion. Then, 100 nm of gold have been deposited using a magnetron sputtering system. Subsequently, a thermal annealing process up to 500°C has been performed to increase the resistor stability over time.

The resistors layout has been obtained by femtosecond laser ablation of the metal layer, using a 10x, 0.25 NA objective, 200 mW of average laser power and 2 mm/s scan velocity. The two resistors have a length of 3 mm along the beam propagation axis, and a width of 15 μm, and they both show an electrical resistance of about 91 Ω. We also implemented some superficial symmetrical trenches, just besides the resistors, to enhance the dynamic response of the device [16]. The trenches are symmetric, 8 μm deep and around 9 μm wide. The final photonic device, with thermal shifters and pins for electrical connections is shown in Fig.1b.

The phase shows a linear dependence with the electric dissipated power, however given the position of the phase shifters on the optical circuit, the applied power at the two shifters influences the phases according to:

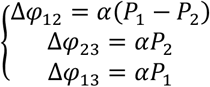

where *α* is a constant, P1 and P2 are the electrical powers applied to thermal shifter 1 and 2 respectively (Fig. 1a). The voltage is related to the power by a quasi-quadratic relation because the electrical resistance is not constant when the temperature increases (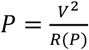, where the resistance *R* is a function of the power *P*). In the ideal case, an increase in *V*_1_ results in an increase of Δ*φ*_13_ and Δ*φ*_12_, but will not affect Δ*φ*_23_. Instead, *V*_2_ rising decreases Δ*φ*_12_, increases Δ*φ*_23_ and leaves Δ*φ*_13_ unchanged. Therefore, a positive error in Δ*φ*_13_ (measured value smaller than expected value) means that *V*_1_ must be increased whereas a positive error in Δ*φ*_23_ is an indicator of a too low *V*_2_. An error in Δ*φ*_12_, on the other hand, is reflecting both on *V*_*1*_ and *V*_*2*_ (See Results and Discussion).

### 3.4 Ferrule fabrication

The technique employed for the ferrule fabrication is called FLICE (femtosecond laser irradiation followed by chemical etching) [19]. This is an established method to realize 3D microstructures in glass materials. It consists in a first fs – laser irradiation to selectively modify the substrate sensitivity to chemical agents, and a consequent chemical bath in an aqueous solution of hydrofluoric acid at 20%, which preferentially removes the irradiated region. We used a 2 mm thick fused silica glass (Foctek) and the same laser used for the direct writing of the integrated photonic circuit. However, for the FLICE process we used the laser second harmonic (515 nm), a 63x/0.75NA objective and an average power of 280 mW. Using these laser parameters, we have irradiated the ferrule, defining the profile along the substrate thickness. We started from the bottom layer of the glass and we have reached the top surface by irradiating multiple concentric planes with an inner separation along the vertical direction of 5 μm. The initial circular radius of the ferrule was 285 μm, but after the chemical etching, the size is reduced to 250 μm.

The advantage of using such a custom-made ferrule is that we can align the PM fibres into an azimuthal polarization, without the need of additional polarizers along the optical path. The polarization configuration is intrinsically reached by the fibres arrangement.

### 3.5 Phase shift

The relative phases between the beams are optimally set when the overall sum of the N patterns results in a homogeneous illumination, to avoid losses of information about the object. The three beams in the illumination produce three carrier frequencies (and their conjugates) in the illumination pattern, each produced by the mutual interference between pairs of the beams. The optimal three relative phase shifts of the *k*-th pattern are given by:

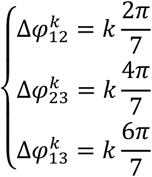

and are indicated in Table 1.

**Table 1.** Optimal relative spatial phases of carrier components in the illumination pattern.

| $k$ | $\Delta\varphi_{12}^k$ | $\Delta\varphi_{23}^k$ | $\Delta\varphi_{13}^k$ |
| --- | --- | --- | --- |
| 0 | 0 | 0 | 0 |
| 1 | $\frac{2\pi}{7}$ | $\frac{4\pi}{7}$ | $\frac{6\pi}{7}$ |
| 2 | $\frac{4\pi}{7}$ | $\frac{8\pi}{7}$ | $\frac{12\pi}{7}$ |
| 3 | $\frac{6\pi}{7}$ | $\frac{12\pi}{7}$ | $\frac{18\pi}{7}$ |
| 4 | $\frac{8\pi}{7}$ | $\frac{2\pi}{7}$ | $\frac{10\pi}{7}$ |
| 5 | $\frac{10\pi}{7}$ | $\frac{6\pi}{7}$ | $\frac{16\pi}{7}$ |
| 6 | $\frac{12\pi}{7}$ | $\frac{10\pi}{7}$ | $\frac{22\pi}{7}$ |

Where the indexes 1, 2 and 3 refer to the pairs of beams coming out from the chip that produce that carrier component. We note that from pattern *k* = 4, we exploited the 2*π* periodicity of the phase to reduce the absolute value of the required phase shift, and consequently of the required electrical power delivered to the phase shifters.

A procedure to measure the actual phase of each carrier component in a single image was developed based on the same technique as used to identify the overall carrier spatial frequency, magnitude and phases as part of the calibration and reconstruction process [3,9]. Here we calculate the cross correlation of a low-pass and a high-pass filtered copy of the image in frequency space. This reveals the carrier components present in the image as peaks in the cross-correlation image whose phases are 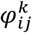. Measuring 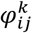 across a set of seven images representing *k* = 0 to 6 lets us compute the measured values of relative phase 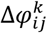 corresponding to the target values shown in Table 1.

### 3.6 SIM reconstruction

The reconstruction of enhanced resolution images from the acquired SIM data follows the method described previously [9], which implements standard algorithms described previously [3] but optimised for processing hexSIM data in a Python environment with GPU acceleration. The procedure is a two-stage process involving a first calibration stage that measures the spatial frequencies, amplitudes and absolute phase of the carrier components in a 7 frame (*k* = 0 to 6) data set. Calibration needs only to be repeated on the timescale of the stability of the illumination system. The second reconstruction phase implements a spatial up-sampling of the data, followed by separation and shifting of the different frequency bands in the image to their correct position given by the spatial frequency of the carriers. Reconstruction also implements low spatial frequency suppression to suppress out-of-focus signals and Wiener filtering to suppress the amplification of noise artefacts.

## 4. Conclusions

We introduced an optical device for the generation of phase-controlled point sources, arranged on a circle, with azimuthal polarization. The chip, installed on the illumination port of a widefield microscope can be used as the light source for structured illumination, upgrading the system to a SIM microscope. Using three sources, an hexSIM pattern can be efficiently formed and dynamically controlling the phase of the chip’s waveguides, we can spatially shift the illumination pattern over the field of view of a widefield microscope. Using this device, we achieved enhanced resolution in imaging biological sample slides.

The present chip opens new possibilities to generate multiple point-sources, with controlled phase. This technology is not only useful for hexSIM but could be used in a variety of configurations [14], maximizing the contrast and the spatial frequency support of SIM reconstructions in epi-illumination microscopy [20]. Along the same line, a higher number of waveguides, controlled in phase and possibly in polarization and amplitude, are conceivable using the present technology, providing the flexibility to generate the patterns for 2D and 3D SIM illumination in different configurations.

## Acknowledgments

The device fabrication was partially performed at PoliFAB, the micro-and nanofabrication facility of Politecnico di Milano (www.polifab.polimi.it). The authors would like to thank the PoliFAB staff for the valuable technical support. Authors acknowledge Dr G. Corrielli for useful discussions on polarization control.

## Funding

European Union under the Horizon 2020 Framework Programme: H2020 Future and Emerging Technologies (801336 -PROCHIP), H2020 Research and Innovation (871124 – Laserlab V).

## Disclosures

MC, PP, FB, RO, MN, HG, AB have filed a patent that relates to some aspects of this work.

## Notes

### Competing Interest Statement

The authors have declared no competing interest.

## References

1. Heintzmann, R., & Huser, T. (2017). Super-resolution structured illumination microscopy. Chemical reviews, 117(23), 13890–13908.

2. Neil, M. A. A., Juškaitis, R., & Wilson, T. (1997). Method of obtaining optical sectioning by using structured light in a conventional microscope. Optics letters, 22(24), 1905–1907.

3. Gustafsson, M. G. (2000). Surpassing the lateral resolution limit by a factor of two using structured illumination microscopy. Journal of microscopy, 198(2), 82–87.

4. Gustafsson, M. G. (2005). Nonlinear structured-illumination microscopy: wide-field fluorescence imaging with theoretically unlimited resolution. Proceedings of the National Academy of Sciences, 102(37), 13081–13086.

5. Jouchet, P., Cabriel, C., Bourg, N., Bardou, M., Poüs, C., Fort, E., & Lévêque-Fort, S. (2021). Nanometric axial localization of single fluorescent molecules with modulated excitation. Nature Photonics, 15(4), 297–304.

6. Chen, B. C., Legant, W. R., Wang, K., Shao, L., Milkie, D. E., Davidson, M. W., … & Betzig, E. (2014). Lattice light-sheet microscopy: imaging molecules to embryos at high spatiotemporal resolution. Science, 346(6208).

7. Reymond, L., Ziegler, J., Knapp, C., Wang, F. C., Huser, T., Ruprecht, V., & Wieser, S. (2019). SIMPLE: Structured illumination based point localization estimator with enhanced precision. Optics express, 27(17), 24578–24590.

8. Förster, R., Lu-Walther, H. W., Jost, A., Kielhorn, M., Wicker, K., & Heintzmann, R. (2014). Simple structured illumination microscope setup with high acquisition speed by using a spatial light modulator. Optics Express, 22(17), 20663–20677.

9. Gong, H., Guo, W., & Neil, M. A. (2021). GPU-accelerated real-time reconstruction in Python of three-dimensional datasets from structured illumination microscopy with hexagonal patterns. Philosophical Transactions of the Royal Society A, 379(2199), 20200162.

10. žurauskas, M., Dobbie, I. M., Parton, R. M., Phillips, M. A., Göhler, A., Davis, I., & Booth, M. J. (2019). IsoSense: frequency enhanced sensorless adaptive optics through structured illumination. Optica, 6(3), 370–379.

11. Van den Eynde, R., Vandenberg, W., Hugelier, S., Bouwens, A., Hofkens, J., Müller, M., & Dedecker, P. (2021). Self-contained and modular structured illumination microscope. Biom Opt Exp 12(7), 4414–4422.

12. Hinsdale, T. A., Stallinga, S., & Rieger, B. (2021). High-speed multicolor structured illumination microscopy using a hexagonal single mode fiber array. Biomedical Optics Express, 12(2), 1181–1194.

13. Pospíšil, J., Wiebusch, G., Fliegel, K., Klíma, M., & Huser, T. (2021). Highly compact and cost-effective 2-beam super-resolution structured illumination microscope based on all-fiber optic components. Optics Express, 29(8), 11833–11844.

14. Ingerman, E. A., London, R. A., Heintzmann, R., & Gustafsson, M. G. L. (2019). Signal, noise and resolution in linear and nonlinear structured-illumination microscopy. Journal of microscopy, 273(1), 3–25.

15. Schropp, M., & Uhl, R. (2014). Two-dimensional structured illumination microscopy. Journal of microscopy, 256(1), 23–36.

16. Calvarese, M., Paiè, P., Ceccarelli, F. Federico Sala, Andrea Bassi, Roberto Osellame & Francesca Bragheri. (2022). Strategies for improved temporal response of glass-based optical switches. Sci Rep 12, 239.

17. Descloux, A., Grußmayer, K. S., & Radenovic, A. (2019). Parameter-free image resolution estimation based on decorrelation analysis. Nature methods, 16(9), 918–924.

18. G. Corrielli, S. Atzeni, S. Piacentini, I. Pitsios, A. Crespi, and R. Osellame. (2018). Symmetric polarization-insensitive directional couplers fabricated by femtosecond laser writing. Opt. Express, 26(12), 15101–15109

19. Osellame, R., Hoekstra, H. J., Cerullo, G., & Pollnau, M. (2011). Femtosecond laser microstructuring: an enabling tool for optofluidic lab-on-chips. Laser & Photonics Reviews, 5(3), 442–463.

20. O’Holleran, K., & Shaw, M. (2012). Polarization effects on contrast in structured illumination microscopy. Optics letters, 37(22), 4603–4605.

